# Somatic Mutations in Vascular Malformations of Hereditary Hemorrhagic Telangiectasia Result in Biallelic Loss of *ENG* or *ACVRL1*

**DOI:** 10.1101/731588

**Authors:** Daniel Snellings, Carol Gallione, Dewi Clark, Nicholas Vozoris, Marie Faughnan, Douglas Marchuk

**Author notes:** Corresponding Author *Email address* (Douglas Marchuk).

## Abstract

Hereditary Hemorrhagic Telangiectasia (HHT) is a Mendelian disease characterized by vascular malformations including visceral arteriovenous malformations and mucosal telangiectasia. HHT is caused by loss-of-function mutations in one of 3 genes; *ENG*, *ACVRL1* or *SMAD4* and is inherited as an autosomal dominant condition. Intriguingly, the constitutional mutation causing HHT is present throughout the body, yet the multiple vascular malformations in HHT patients occur focally, rather than manifesting as a systemic vascular defect. This disconnect between genotype and phenotype suggests that a local event is necessary for the development of vascular malformations. We investigated the hypothesis that local somatic mutations seed the formation HHT-related telangiectasia in a genetic two-hit mechanism. We identified low-frequency somatic mutations in 9/19 telangiectasia using high depth next-generation sequencing. We established phase for 7 of 9 samples using long-read sequencing, which confirm that the germline and somatic mutations in all 7 samples exist in trans configuration; consistent with a genetic two-hit mechanism. These combined data suggest that biallelic loss of *ENG* or *ACVRL1* may be a required event in the development of telangiectasia, and that rather than haploinsufficiency, vascular malformations in HHT are caused by a Knudsonian two-hit mechanism.

## Introduction

Hereditary Hemorrhagic Telangiectasia (HHT) is a Mendelian disease characterized by the development of multiple focal vascular malformations consisting of arteriovenous malformations in visceral organs and telangiectasia in mucosal and cutaneous tissue. The genetic etiology of HHT has been established and is caused by mutations in *ENG* ^1^, *ACVRL1* ^2^, and rarely *SMAD4* ^3^; all of which follow an autosomal dominant inheritance pattern. Despite our understanding of the genetics of and downstream pathways involved in HHT, the molecular mechanisms that initiate HHT-related vascular malformation are poorly understood. Early studies of the functional consequences of HHT causal mutations established that these result in the loss of function of the gene product. These findings, in combination with autosomal dominant inheritance, led to the presumption that vascular malformations result from haploinsufficiency of the mutated gene product.^4–6^ However, haploinsufficiency does not account for why HHT-related vascular malformations occur as strictly focal lesions, despite the systemic presence of the causal germline mutation. This disconnect between genotype and phenotype led to an alternative long-standing hypothesis that HHT-related vascular malformations result from a Knudsonian two-hit mechanism, where a local somatic mutation in the wild type allele of the causal gene seeds the formation of focal lesions.

The only published study which directly addresses the two-hit hypothesis attempted to determine whether Endoglin was present on the endothelial lining of arteriovenous malformations from an HHT patient with a causal mutation in the corresponding gene *ENG*. ^7^ Endoglin immunostaining was visible in the vessel lining, albeit at low levels. The presence of Endoglin in HHT-associated vascular malformations would contradict a two-hit mechanism, however complete loss of staining might not be predicted to occur, especially with the heterogeneous – and potentially mosaic – tissue of an arteriovenous malformation that may only contain a minority of cells that harbor the somatic mutation. Previous attempts to address this hypothesis at the DNA level have been hampered by the limitations of past sequencing technology. The advent of next-generation sequencing has drastically increased our sensitivity for detecting low-frequency somatic mutations. Somatic mutations have been identified in a diverse array of vascular malformations ^8–14^ including recent evidence that sporadic arteriovenous malformations, not associated with HHT, harbor somatic activating mutations in *KRAS* or *MAP2K1*. ^15,16^ Notably, a genetic two-hit mechanism is known to contribute to Cerebral Cavernous Malformations (CCM) ^17,18^ and Capillary Malformation – Arteriovenous Malformation Syndrome (CM-AVM) ^19^; like HHT, both diseases are caused by autosomal dominant loss of function mutations. Here we demonstrate that HHT-related telangiectasia contain biallelic mutations in *ENG* or *ACVRL1*, resulting in homozygous loss of function; evidence in support of the long-standing hypothesis that telangiectasia pathogenesis follows a genetic two-hit mechanism.

## Materials and Methods

### Sample Collection

Individuals were enrolled in the study after giving informed consent (approved by either the St. Michael’s Hospital IRB committee or the Duke University Health System IRB Committee). Diagnosis of HHT was based on identification of a pathogenic germline mutation, or by exhibiting at least three of the four symptoms as per the Curaҫao criteria (Supp Table 1). ^20^

Telangiectasia were resected using a 3mm punch biopsy, after local anesthesia (1% xylocaine with epinephrine), with standard aseptic technique. Samples 6005-1 was immediately formalin fixed (10% formalin) and paraffin embedded (FFPE), and then shipped at room temperature. All other samples were immediately frozen at −80 Celsius, and then shipped on dry ice. Saliva samples were obtained using Oragene DNA saliva kits at the time of tissue collection. Blood from patient 6003 was obtained during a subsequent visit, shipped at room temperature, and immediately used for RNA extraction.

### DNA/RNA Extraction

DNA from telangiectasia samples was extracted using the DNeasy Blood and Tissue Kit (Qiagen). DNA from FFPE sample 6005-1 was extracted using QIAamp DNA FFPE Tissue Kit (Qiagen). Genomic DNA and RNA were extracted from peripheral blood leukocytes from patient 6003 and from a non-HHT control individual using the Gentra PureGene Blood Kit (Qiagen) and TRIzol Reagent (Invitrogen) extraction protocols, respectively, as per the manufacturers’ directions. Nucleotides are numbered with the conventional nomenclature of the A in the first ATG as c.1.

### Targeted Sequencing

To enable the detection of somatic mutations in telangiectasia we used a high-depth next generation sequencing strategy. Somatic mutations involved in the pathogenesis of other vascular malformation diseases such as cerebral cavernous malformations often have a low allele frequency due to somatic mosaicism in the malformation. Somatic mosaicism is also present in retinal AVMs from mouse models of HHT suggesting that low allele frequency may be a confounder when identifying somatic variants in telangiectasia. In addition, the telangiectasia samples collected for this study consist of bulk biopsied tissue which have not been enriched for any particular cell type. Considering these potential sources of normal (non-mutant) cell contamination, we sequenced telangiectasia to >1000x coverage and incorporated a unique molecular identifier to enable the detection of variants as low as 0.1% allele frequency.

Eighteen fresh-frozen samples and one FFPE sample was sequenced using a custom Agilent SureSelect panel covering 16 genes implicated in various vascular malformation disorders: *ENG*, *ACVRL1*, *SMAD4*, *BRAF*, *CCM2*, *FLT1*, *FLT4*, *GNAQ*, *KDR*, *KRAS*, *KRIT1*, *MAP2K1*, *NRAS*, *PDCD10*, *PIK3CA*, and *PTEN*. To ensure the generation of high-quality sequencing libraries, the Agilent NGS FFPE QC Kit was used to determine the extent of DNA degradation in the FFPE sample. Samples with ∆∆Cq values >2 were excluded from the study as per manufacturer recommendation. Sequencing libraries were generated using the Agilent SureSelect XT HS Kit. Samples were then pooled and sequenced on an iSeq 100 (Illumina) with paired-end 150bp reads. All samples were sequenced to a mean depth of >1000x.

### Mutation Detection

Sequencing data was processed and analyzed using a custom pipeline based on the GATK best practices for somatic short variant discovery. Briefly, after analyzing the raw data with fastQC to ensure high quality data, the adapter sequences were trimmed from reads using bbduk, reads were aligned to the hg19 human reference genome using bowtie2, duplicates were removed based on UMI sequence using fgbio, variants were called using MuTect2 in tumor-only mode, and variants were annotated using snpSift. The resulting variant call file (VCF) was filtered through several steps to identify somatic mutations. To identify variants that may change the protein sequence or impact splicing, we selected for variants that occur within exons or within 10bp of an exon. From this set, we removed variants that are present in the population at >0.01% frequency by comparing to 3 databases (dbSNP, 1000 Genomes project, and the Exome Aggregation Consortium [ExAC]); as these variants are more common than the frequency of HHT (1-5 in 10000). ^21^ We removed any variants present in <0.02% of sequence reads, as this is the reported technical limit of detection for the SureSelect XT technology. We also removed variants in regions with <100x coverage, variants with <5 supporting reads, variants that were strand specific, and variants where <50% of alternative bases had a quality score >30 (>Q30).

Candidate somatic variants identified in the targeted sequencing data were then validated by sequencing amplicons generated during a second, independent round of PCR amplification. We designed primers for each sample to specifically amplify the position of the somatic mutation and 100-200bp of flanking sequence. When possible, the primers were designed such that they would capture the position of both the germline and somatic mutations within a single amplicon. Each primer was synthesized with the following Illumina flow cell adapter sequences such that the amplicons could be easily indexed and sequenced: Forward 5’-TCGTCGGCAGCGTCAGATGTGTATAAGAGACAG-[sequence-specific primer]-3’, Reverse 5’-GTCTCGTGGGCTCGGAGATGTGTATAAGAGACAG-[sequence-specific primer]-3’. Amplicons were prepared from telangiectasia DNA and constitutional DNA if available (Supp Table 1) using two rounds of PCR. The first round of PCR was 25 cycles and served to amplify the target region from genomic DNA. Amplicons from the first round were purified using AMPure XP beads (Beckman Coulter) and used for a second round of PCR using 8 cycles to attach a sample index using the Nextera XT Index Kit. Amplicons from the second round were purified, pooled, and sequenced on an iSeq 100 with 150bp paired-end reads to a depth of >10000x. The frequency of the somatic mutation in telangiectasia and constitutional DNA was determined using custom scripts, excluding bases <Q15.

As these amplicons were sequenced in the same run and at high depth, it is possible for a low level of index misassignment causing switching of reads between samples. This could cause some reads from telangiectasia to be assigned as constitutional reads and vice versa. To estimate the rate of index misassignment, we examined pairs of samples that target different genomic locations and quantified the proportion of misassigned reads between these samples. Based on this we estimate that the rate of misassignment is 0.2 – 0.8%, relative to the sample of origin. For example, if one sample has a somatic mutation with a frequency of 1% and was sequenced to 100000x coverage, then 2 – 8 reads containing the somatic mutation would be misassigned to each of the other samples in the pool.

### Establishing Phase

To establish the phase of the somatic and germline mutations we used either short-read sequencing with Illumina chemistry, or long-read sequencing with PacBio chemistry depending on the distance between the two mutations. For mutations <500bp apart, we generated amplicons during the validation process that cover the positions of both the somatic and germline mutations. These amplicons were sequenced on an iSeq 100 as described above.

For mutations >500bp we designed primers such that the resulting amplicon would cover the position of both the germline and somatic mutations for each sample. Each primer was synthesized with the PacBio ‘universal tag’ as follows to enable indexing and sequencing: Forward 5’-/5AmMC6/ GCAGTCGAACATGTAGCTGACTCAGGTCAC-[sequence-specific primer]-3’, Reverse 5’- /5AmMC6/TGGATCACTTGTGCAAGCATCACATCGTAG-[sequence-specific primer]-3’. We generated amplicons spanning the somatic and germline mutations using the LongAmp Taq DNA Polymerase Kit as per manufacturer instructions. These amplicons were purified with AMPure XP beads and used for a second round of PCR to attach the sample index. This process used no more than 30 cycles of PCR total. These amplicons were pooled and sequenced across one SMRT cell on a PacBio Sequel System. The sequence reads were aligned to the hg19 human genome using Minimap2 in ava-pb mode.

The single molecule resolution of these technologies allowed us to determine how the mutant alleles are arranged; if the mutations are in trans then reads will have either the somatic mutant allele or the germline mutant allele, if the mutations are in cis then reads will have either no mutant alleles or both mutant alleles. In total we generated mutation-spanning reads for 7 telangiectasia, each with >100 reads which contained the somatic mutation. The p-values reported for phase status were calculated using a binomial distribution with the null hypothesis that a random mutation has an equal probability of cis or trans configuration with a nearby variant.

The genomic distance between mutations varied greatly with the closest mutations in sample 6005-1 with 26 bases between mutations, and the most distant in 6001-10 with 18.7 kilobases between mutations. Of the 9 telangiectasia with identified somatic mutations, the distance between mutations in 6002-2, 6003-1, and 6005-1 was small enough to allow for mutation-spanning reads using illumina chemistry (Table. 2). Mutation-spanning reads were generated for 6001-3, 6001-7, 6001-8, and 6002-1 using PacBio chemistry. We were unable to generate amplicons spanning the mutations for 6001-1 and 6001-10. The sequence downstream of the somatic mutation in these telangiectasia contains several repetitive regions which, combined with the genomic distance and limited quantity of input DNA may have contributed to PCR failure.

One notable confounder in this analysis is the generation of chimeric reads resulting from template switching during PCR. The generation of chimeric reads is known to interfere with amplicon-based haplotype phasing by switching a variant from one strand to another, potentially generating new haplotypes not present in the original sample. ^22^ In practice, chimeric reads randomize the arrangement of the somatic and germline mutations. The frequency of chimeric arrangements is highly dependent on the distance between mutations and the number of PCR cycles used to make the amplicons. To reduce the number of chimeric reads in our libraries we used no more than 30 cycles for amplification. A previous study reports a chimeric arrangement frequency of 6.5% for 29 cycles of amplification for mutations 9kbp apart. ^22^ Chimeric reads may account for the very few discordant reads in some of our samples, as shown in Table 2. Nonetheless, these were so minor in comparison to the great majority of the reads that phase could be unequivocally determined.

### in vitro Splicing

A 3.8kb fragment of *ACVRL1* genomic DNA spanning from 431 bases upstream of exon 3 to 215 bases downstream from exon 8 was amplified and this entire insert was ligated into the MCS of pSPL3, a splicing vector. ^23^ Clones were sequenced to ensure that no PCR-induced errors were present in the exons and adjacent intronic regions of the insert. The specific mutation, c.625+4A>T, was introduced using site directed mutagenesis and again, clones were sequenced to verify that the only sequence difference was at the intended site. Plasmid DNA from empty vector, wild type (control) vector and mutation-containing vector were transfected into HEK293T cells using Lipofectamine 3000 (ThermoFisher Scientific), incubated for 24 hours, and then the RNA was extracted using TRIzol and Direct-zol RNA miniprep kit (Zymo Research).

### Reverse-Transcription PCR

RNA extracted from patients’ blood and from transfected cells was used as template for cDNA synthesis using the Maxima H Minus First Strand cDNA kit (ThermoFisher Scientific). An RT primer in *ACVRL1* exon 8 was used in the RNA from patients’ blood while a vector-specific RT primer was used for the RNA from transfected cells to ensure that only RNA from the transfected vectors was being used as template for cDNA synthesis. cDNA from the patients was PCR amplified using primers in exons 3 and 6 while cDNA from the transfected cells was PCR amplified using primers in exons 3 and 8. PCR reactions were run on 1% agarose gels, the bands excised and Sanger sequenced.

## Results

To determine whether a genetic two-hit mechanism underlies HHT pathogenesis, we tested three underlying expectations of the two-hit mechanism: 1) telangiectasia contain a somatic mutation in the same gene as a germline mutation which causes HHT, 2) the somatic and germline mutations are biallelic, and 3) both mutations result in loss-of-function.

### Telangiectasia Harbor a Somatic Mutation in ENG or ACVRL1

We used capture-based library preparations to sequence 19 telangiectasia for the three HHT causal genes (*ENG*, *ACVRL1*, and *SMAD4*) and 13 other vascular malformation-related genes (see Methods for identity of genes). The 13 non-HHT vascular malformation genes were chosen with the possibility that they also might harbor somatic mutations, but primarily to serve as control genes, since the two-hit mechanism requires a mutation in the corresponding HHT gene harboring the causal germline mutation. Somatic mutations in these other genes may or may not contribute to HHT pathogenesis, but absence of a somatic mutation in the casual HHT gene, would violate the first expectation of the genetic two-hit mechanism.

In each telangiectasia we identified a pathogenic germline mutation in either *ENG* or *ACVRL1*. Although in most cases the patient’s germline mutation was already known from clinical diagnostic sequencing, we intentionally remained blinded to this information until after our own sequence analysis of the tissue samples. In 6003-1, the patient harbors a silent germline mutation which was found by the clinical lab and noted as a variant of unknown significance. Below we show that this variant is indeed the pathogenic germline variant in this patient.

We used the MuTect2 variant caller to detect variants present in the sequence data. To identify candidate somatic mutations, we removed variants based on several stringent filtering criteria including briefly; intronic or intergenic variants, population frequency >0.01%, <0.1% or <5 total supporting reads, low coverage, strand specificity, and low base quality scores.

To validate or refute the authenticity of each candidate somatic mutation we performed an independent round of amplification using primers flanking each putative variant position for each sample, and sequenced to >10000x coverage. In each tissue sample, the identical somatic variant was re-identified. Thus, these variants were bona fide somatic mutations existing in the telangiectatic tissue. In total, we identified somatic variants in 9 of 19 telangiectasia; 5 in *ENG*(NM 001114753) 4 in *ACVRL1*(NM 000020) (Table 1) (Fig. 1) (See Methods). In each case, the somatic mutation was found in the same gene as the pathogenic germline mutation. Somatic mutations were not found in any of the other 15 genes sequenced, not even in one of the other HHT casual genes. Importantly, no telangiectasia harbored more than a single somatic mutation. The lack of mutational noise suggests these mutations are pathobiologically significant. Importantly, all are consistent with strong mutations; five of the variants are small indels resulting in a frameshift, three are in-frame indels, and one is a point mutation 4 bases after an exon-intron boundary that is predicted to impact RNA splicing.

**Figure 1:**
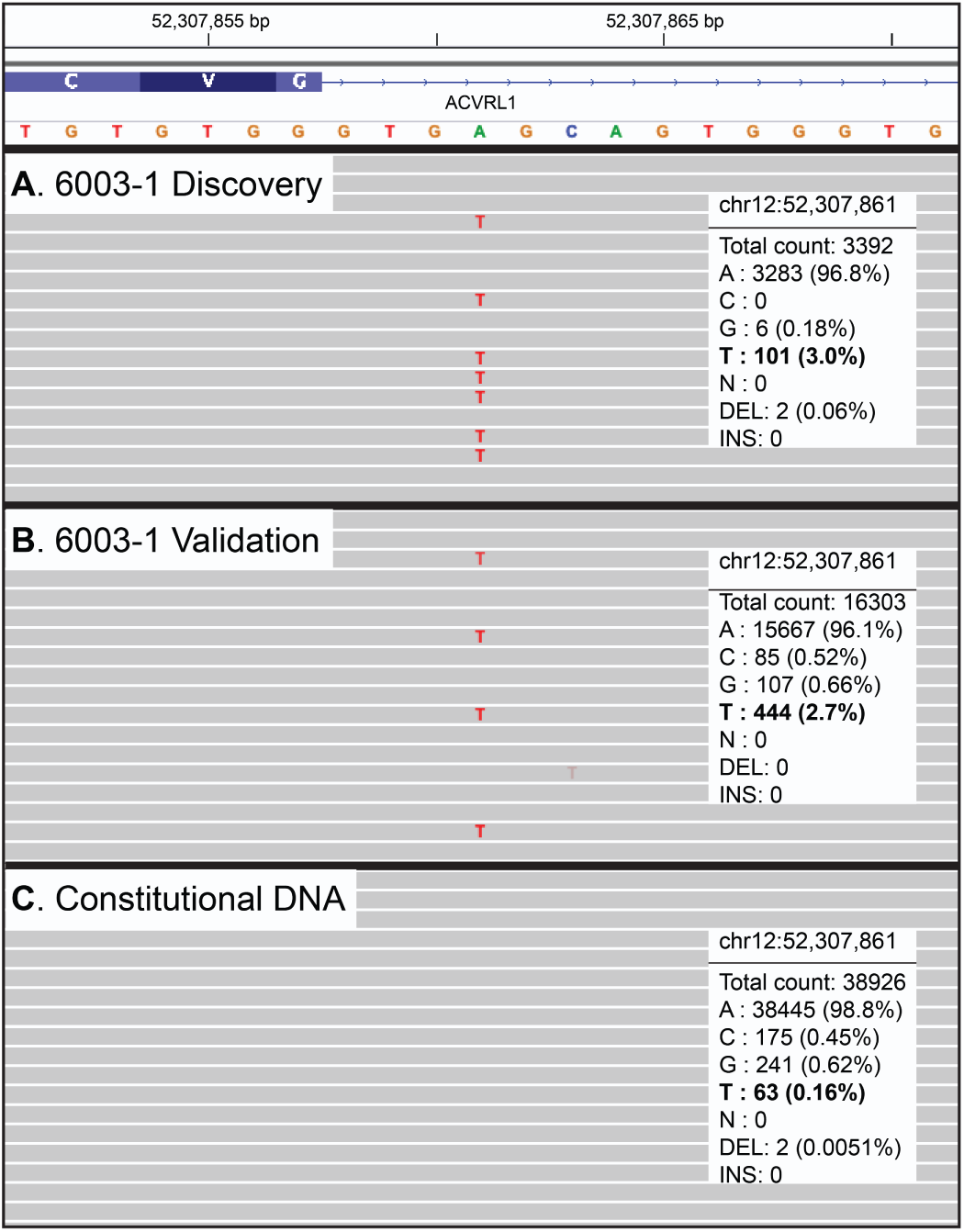
Low Frequency Somatic Mutations Detected in Telangiectasia. Visualization with IGV of deep next-generation sequencing data for one representative sample with a somatic mutation in *ACVRL1*. The somatic mutation in 6003-1, an A>T point mutation, is present at low frequency in DNA from telangiectatic tissue used for (A) discovery (capture-based sequencing) and (B) validation (amplicon-based sequencing). The mutation is below the level of sequencing noise in (C) constitutional DNA (amplicon-based sequencing) confirming the mutation is somatic.

**Table 1.**
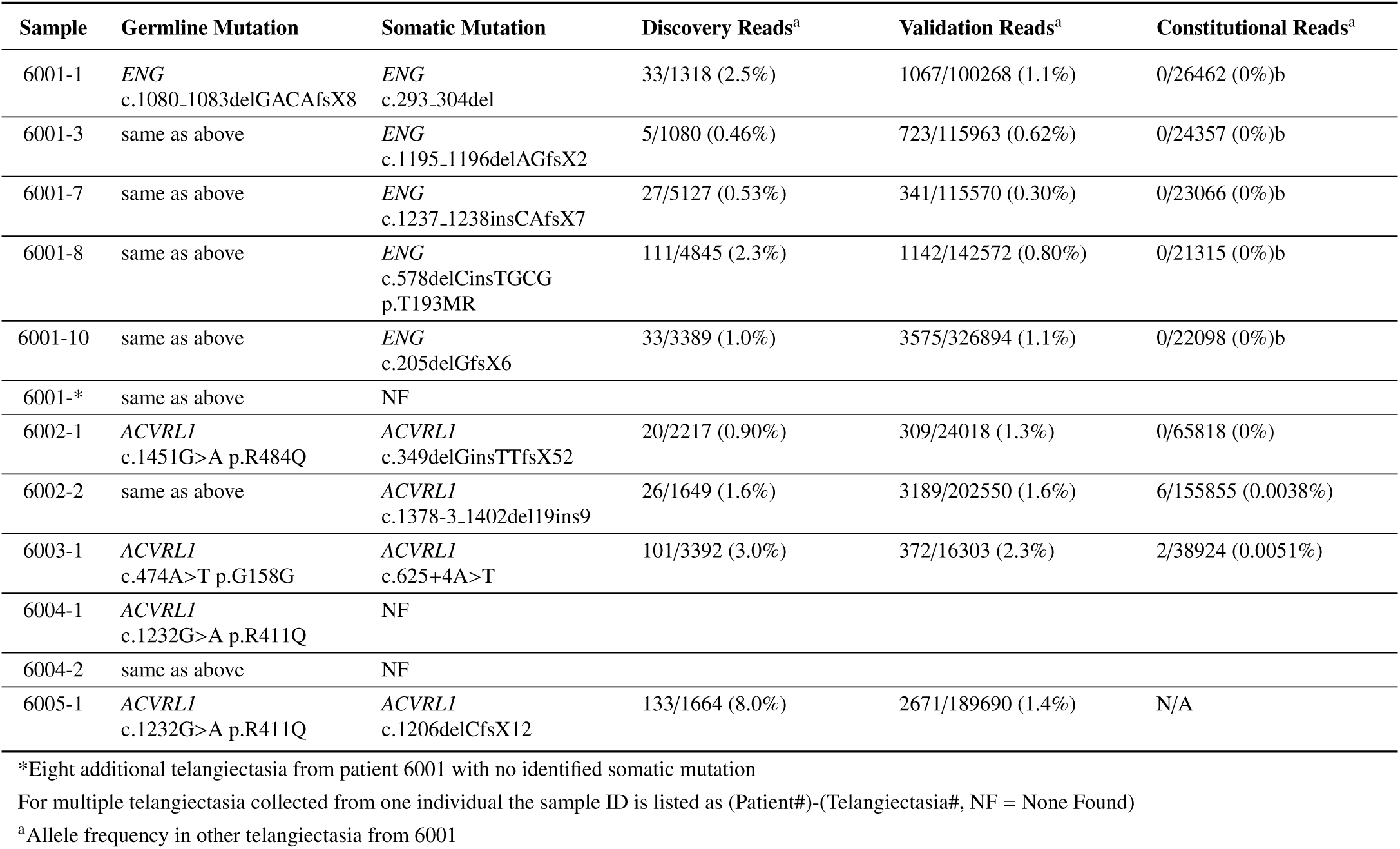
Summary of Somatic Mutation Discovery and Validation Sequencing

Although these variants fall well below the 50% allele frequency expected for germline variants, it is formally possible that these variants exist constitutionally as very rare, somatic mosaic variation in the patients. We next investigated whether these somatic mutations were present in constitutional DNA from the patient. A source of constitutional DNA (buccal epithelial cells from saliva) was available for seven of the nine mutation-positive samples, and in each we find that the somatic mutation was completely absent or present at a level no higher than technical sequencing noise in that sample (see Methods). Saliva was not available for patient 6001, but we obtained and sequenced DNA from multiple telangiectasia collected from this same patient. This enabled us to determine whether any of the five somatic mutations that we identified in individual samples was present in tissue of near identical pathobiology from the same individual; compared with saliva as a control, this is a more powerful test for somatic mosaicism. We found that the somatic mutations identified in five of the telangiectasia for which we identified a mutation were entirely absent in all other telangiectasia from this same individual. Finally, Sample 6005-1 is a single archived FFPE telangiectasia and no source of constitutional DNA is available. In total, we found that 9 of 19 telangiectasia harbor a somatic mutation specifically in the same gene as a pathogenic germline mutation and that these mutations are not present constitutionally. The presence of somatic mutations in telangiectasia fulfills the 1st expectation of the genetic two-hit mechanism.

### Somatic and Germline Mutations are Biallelic

The 2nd expectation of the genetic two-hit mechanism is that the somatic and germline mutations are biallelic; such that the somatic mutation occurs on the wild-type allele of the causal gene, in trans with the germline mutation. To determine if the mutations are biallelic we examined whether they were arranged in a cis or trans configuration by sequencing amplicons that cover the nucleotide positions of both somatic and germline mutations in a single molecule. The amplicons were sequenced with either short-read (Illumina) or long-read (PacBio) chemistry, depending on the amplicon size, in order to generate reads that would span the two mutations. In contrast to traditional Sanger sequencing which measures the population average at each position, both Illumina and PacBio chemistries output sequences of single DNA molecules. In total we generated mutation-spanning reads for 7 telangiectasia, each with more than 100 reads that contained the somatic mutation. From these mutation-spanning reads we established that >95% of reads with the somatic mutation possessed the wild type allele at the position of the germline mutation, showing that all 7 mutation pairs are in trans configuration (Table 2) (Fig. 2A-B). Any two variants in a chromosome must be arranged in cis or trans with an equal probability of either arrangement. Considering this, our observation that 7/7 mutation pairs are arranged in trans corresponds to a p-value of 0.008 demonstrating significant bias towards a trans configuration. These data show that the somatic and germline mutations are biallelic, fulfilling the 2nd expectation of the genetic two-hit mechanism.

**Figure 2:**
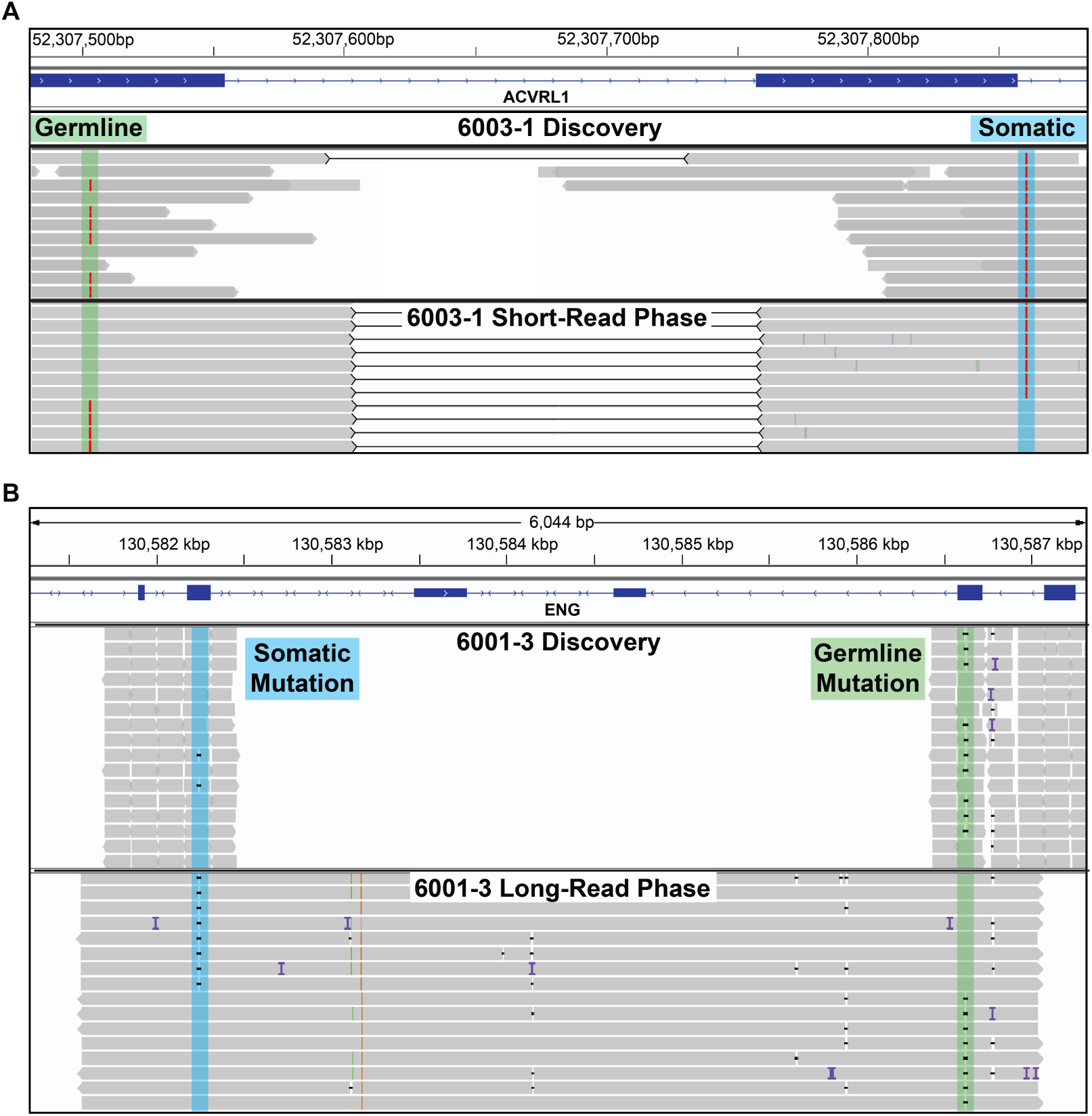
Somatic and Germline Mutations are Biallelic. IGV visualization of two samples showing both methods of establishing phase. Each panel shows reads from the initial discovery sequencing and the reads used to establish phase. (A) Somatic and germline mutations in 6003-1 are both A>T point mutations and highlighted by the blue and green regions respectively. Since the distance between these mutations is relatively small (357bp) phase was established using Illumina short-reads (also used for validation). Black lines between reads denote read pairs, showing that both reads originate from a single molecule of DNA. Each molecule with the somatic mutation contains the wild-type allele at the germline mutation position proving the mutations are biallelic. (B) Somatic and germline mutations in 6001-3, both small deletions. The genomic distance between these mutations is 4377bp. In long reads that span the two mutations, each read with the somatic mutation contains the wild-type allele at the position of the germline mutation.

**Table 2.** Phase of Somatic and Germline Mutation Pairs

| Sample | Total Reads | Trans Reads | Cis Reads | P-value |
| --- | --- | --- | --- | --- |
| 6001-1 | N/A | N/A | N/A | N/A |
| 6001-3 | 112 | 112 (100%)<sup>a</sup> | 0 (0%) | 1.9e-34 |
| 6001-7 | 155 | 153 (98.7%)<sup>a</sup> | 2 (1.3%) | 2.6e-43 |
| 6001-8 | 593 | 590 (99.5%)<sup>a</sup> | 3 (0.5%) | 1.0e-171 |
| 6001-10 | N/A | N/A | N/A | N/A |
| 6002-1 | 125 | 120 (96.0%)<sup>a</sup> | 5 (4.0%) | 5.5e-30 |
| 6002-2 | 3189 | 3160 (99.0%)<sup>bc</sup> | 29 (1.0%) | 4.2e-890 |
| 6003-1 | 372 | 364 (97.8%)<sup>bc</sup> | 6 (1.4%) | 1.4e-99 |
| 6005-1 | 2671 | 2653 (99.3%)<sup>bc</sup> | 18 (0.7%) | 6.3e-759 |

<sup>a</sup>Reads generated with PacBio long-read chemistry)
<sup>b</sup>Reads generated with Illumina short-read chemistry
<sup>c</sup>These reads were also used for validation shown in Table 1

### Mutations are Consistent with Homozygous Loss of Function

The 3rd expectation of the genetic two-hit mechanism is that the biallelic somatic and germline variants both result in loss of function. Due to the functional studies and extensive allelic series of mutations in each of the HHT genes, HHT is known to be caused by loss of function mutations. The germline mutation in 3 of the 5 patients in this study has been identified previously in an individual with HHT and are reported in ClinVar (6001:VCV000213214.2, 6002:VCV000212796.2, 6005:VCV000008243.2). There are also several publications supporting the pathogenicity of these mutations. ^24–28^ These are all therefore bona fide loss of function mutations. The germline mutation in 6003-1, a silent mutation in *ACVRL1* exon 4, has been identified before in an individual with HHT, however it was classified as a variant of unknown significance (VUS). We used the in silico tool Human Splicing Finder 3.1^29^ to analyze this variant and found that it was predicted to both disrupt an exonic splice enhancer and create an internal splice donor site, potentially activating a cryptic splice site. Based on this prediction, we extracted RNA from peripheral blood leukocytes of patient 6003 and a control individual and performed RT-PCR to examine the splicing of *ACVRL1* transcripts. RNA from patient 6003 shows a new splice variant that is not present in control wild-type RNA (Fig. 3B). As predicted by the Splice Finder program, the aberrant transcript is spliced precisely at the internal splice donor created by the mutation. The resulting transcript is missing the portion of exon 4 downstream of the germline mutation and skips exon 5 resulting in the in-frame deletion of 52 amino acids (Fig. 3D-E). This deleted region contains several codons with known pathogenic missense mutations, suggesting that the 52 amino acid deletion would likely also result in loss of function. It is possible that the skipping of exon 5 is due to alternative splicing observed only in peripheral blood leukocytes, rather than a result of the mutation. If exon 5 is retained, the mutation would then generate a protein lacking 17 amino acid residues from exon 4 but then be frameshifted for the remainder of the transcript. With this evidence, all of the identified germline mutations meet the American College of Medical Genetics (ACMG) criteria for pathogenic mutations ^30^, fulfilling the first half of the 3rd expectation of the genetic two-hit mechanism.

**Figure 3:**
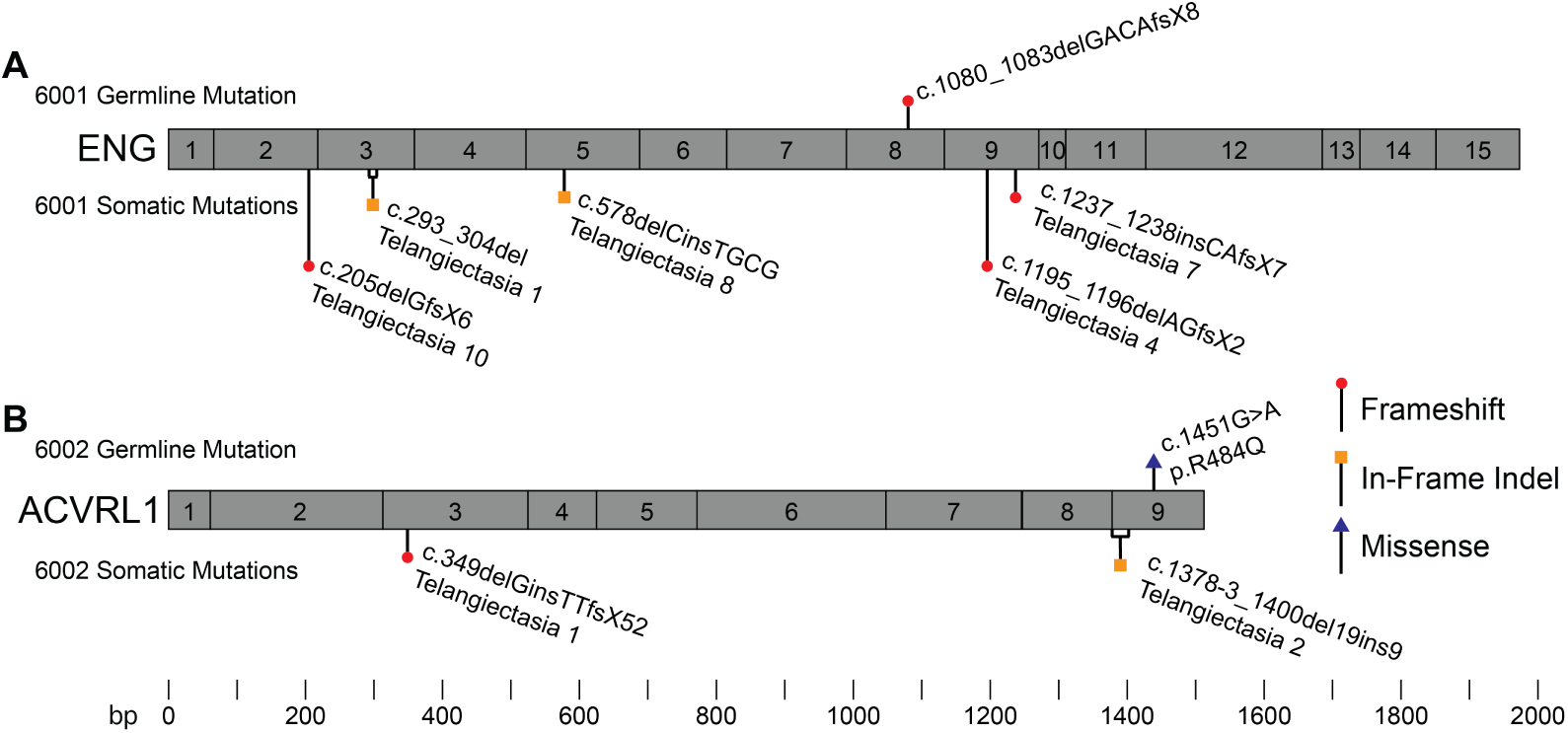
Multiple Telangiectasia from Single Individuals Contain Unique Somatic Mutations. Schematic representation of exons in *ENG* and *ACVRL1* with germline and somatic mutations identified in (A) 5 telangiectasia collected from patient 6001 and (B) 2 telangiectasia collected from patient 6002. In each panel the common germline mutation is listed above the gene and somatic mutations in each telangiectasia below the gene. Gene structure and mutation position are to scale.

In contrast to the germline mutation, many of which have been previously identified, the somatic mutations we identified are all novel. The ACMG guidelines for establishing pathogenicity are not applicable to somatic variants, however several lines of evidence support that all of the somatic mutations result in loss of function. Five of the nine somatic mutations result in a frameshift; all resulting in premature termination codons which would generate transcripts susceptible to nonsense-mediated decay. Frameshift mutations in *ENG* or *ACVRL1* are the most common mechanism for loss of function leading to HHT. Based on this, the 5 somatic frameshift mutations likely result in loss of function. Other than frameshift mutations, the other 4 somatic mutations we identified consisted of 3 in-frame deletions and 1 intronic mutation predicted to impact splicing. These 4 mutations are not present in the genome aggregation database (gnomAD) showing that the population allele frequency of these variants is extremely low or zero. For the somatic in-frame deletion mutation found in 6001-8, there are two reports of different in-frame deletions with overlap at this position which are known to cause HHT, suggesting that the somatic deletion in 6001-8 is likely to result in loss of function. The somatic mutations in 6001-1 and 6002-2 also result in inframe deletions, which delete 4 and 7 amino acids respectively. Comparing the crystal structures of *ENG* and *ACVRL1* we determined that the somatic mutations in 6001-1 and 6002-2 delete portions of a beta strand and helix respectively, potentially impacting protein folding (Table 3). The in silico tool PROVEAN was used to predict how the protein would tolerate these deletions. The threshold of −2.5 or lower (more negative) is considered a deleterious change. The scores for 6001-8, 6001-1, and 6002-2 were −6.106, −14.903, and −25.903 respectively, strongly suggesting that all three are deleterious. The remaining somatic mutation is intronic, and occurs 4 nucleotides from the exon-intron boundary. We used Human Splicing Finder 3.1 to predict the effect of this variant on splicing, and found that it is likely to disrupt the acceptor site. This prediction was confirmed by RT-PCR using an in vitro splicing construct which revealed that the somatic mutation prevents the formation of full-length *ACVRL1* transcripts (Fig. 3C). In summary we present evidence supporting that the biallelic germline and somatic mutations all likely result in loss of function, fulfilling the 3rd expectation of the genetic two-hit mechanism.

**Table 3.** Evidence that Identified Germline and Somatic Mutations Result in Loss of Function

| Sample | Germline Mutation | Somatic Mutation |
| --- | --- | --- |
| 6001-1 | Frameshift: ACMG PVS1<br>6 supporting publications | In-Frame Deletion (-4 residues)<br>PROVEAN: Deleterious (-14.903)<br>deletes region in β-sheet<br>gnomAD AF: 0 |
| 6001-3 | same as above | Frameshift<br>common ENG LOF mechanism, expect NMD |
| 6001-7 | same as above | Frameshift<br>common ENG LOF mechanism, expect NMD |
| 6001-8 | same as above | In-Frame Delins (-1 +2 residues)<br>2 pathogenic in-frame indels overlapping this codon<sup>31,32</sup><br>PROVEAN: Deleterious (-6.106)<br>gnomAD AF: 0 |
| 6001-10 | same as above | Frameshift<br>common ENG LOF mechanism, expect NMD |
| 6002-2 | same as above | In-Frame Delins (-7 +1 residues)<br>PROVEAN: Deleterious (-25.903)<br>deletes region in α-helix<br>gnomAD AF: 0 |
| 6003-1 | Cryptic Splice Site: ACMG PS3<br>in silico predicted to activate<br>cryptic site<br>in vitro evidence (Fig. 4) | Splice Site<br>in silico predicted to disrupt donor site<br>gnomAD AF: 0 |
| 6005-1 | Missense: ACMG PS1<br>16 supporting publications | Frameshift<br>common ACVRL1 LOF mechanism, expect NMD |

AF = Allele Frequency; NMD = Nonsense Mediated Decay
Germline variant classification according to ACMG guidelines, not applicable to somatic variants<sup>30</sup>
PVS1 = Very strong evidence for pathogenicity; PS1-4 = Strong evidence
PROVEAN scores below -2.5 are predicted deleterious

### Telangiectasia from the Same Individual Harbor Unique Somatic Mutations

We next sought to determine whether mutant cells in different telangiectasia derive from a somatic mutation in a common ancestor cell, or whether the mutant cell population in each telangiectasia derives from an independent somatic mutation event. To test this, we examined the somatic mutations present in multiple telangiectasia from single individuals. In patient 6001, for which we had obtained 13 different telangiectasia, we identified a somatic mutation in 5 telangiectasia tissue samples. In each case the somatic mutation in each telangiectasia was unique. Likewise, for patient 6002, for which we had two telangiectasia, we identified a unique somatic mutation in each (Fig. 4). These results are consistent with independent mutation events rather than the somatic mutation occurring in a progenitor cell or clonality due to a metastasis from a single initial lesion.

**Figure 4:**
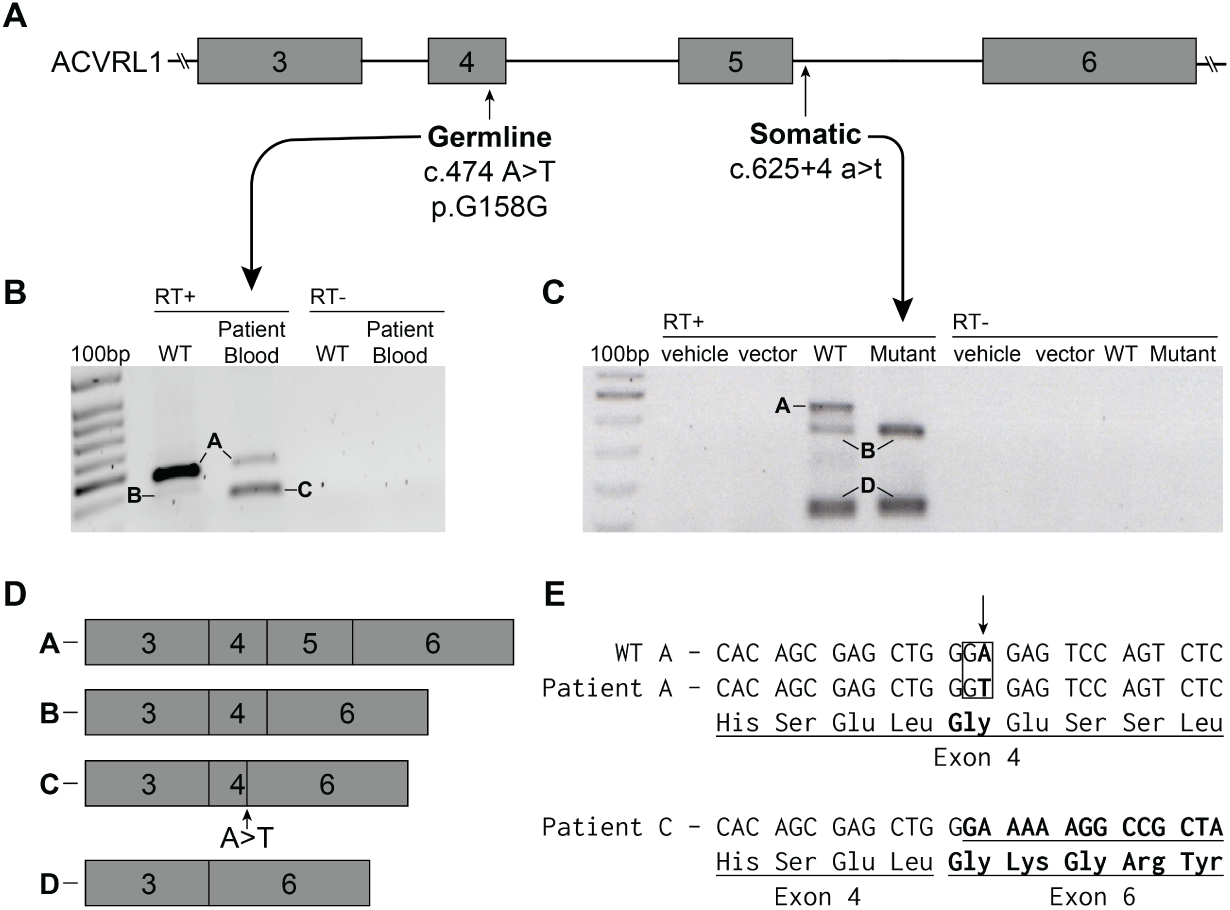
Germline and Somatic Mutations in 6003-1 Both Cause Aberrant Splicing of ACVRL1 RNA. (A) The gene structure of *ACVRL1* exons 3-6 marked with the location of the germline and somatic mutations found in 6003-1. (B) RT-PCR showing *ACVRL1* transcripts from blood leukocytes taken from wild-type (WT) and patient blood containing the germline mutation. The labeled bands were excised and sequenced. Full length transcript (Band A) is present in both the control and the patient although the level in the patient is greatly reduced. Band B in the control contains complete exons 3, 4 and 6. This splice variant has been seen previously and differs from band C in the patient which splices from the newly created splice acceptor site within exon 4 directly to exon 6. (C) As the somatic mutation is only present in 3.0% of reads it would be challenging to detect mis-spliced RNA from the biopsied tissue. Therefore, we inserted wild-type and mutant sequence of *ACVRL1* into an in vitro splicing vector, pSPL3-ACVRL1, and used RT-PCR to visualize the impact of the mutation on splicing. Only WT band A shows the full-length transcript containing exon 5; there is no corresponding full-length transcript from the plasmid containing the somatic mutation. (D) Exon structure of *ACVRL1* transcripts determined by sequencing the excised bands. (E) Sequence of DNA showing the nature of the germline mutation. In *ACVRL1* transcripts containing the germline mutation, exon 4 is shortened due to the activation of a cryptic splice site.

## Disscussion

In this study we present strong evidence that vascular malformations associated with HHT, specifically, cutaneous telangiectasia, follow a genetic two-hit mechanism of pathogenesis. HHT is also associated with arteriovenous malformations in lung, liver, brain and the gastrointestinal tract, but these tissues are not available for prospective collection. We postulate that the visceral, deeper vascular malformations that occur in HHT also follow this two-hit mechanism.

The two-hit hypothesis for HHT pathogenesis has persisted for decades without evidence, but these low frequency somatic mutations are challenging to identify using traditional sequencing methods. The only published study to address this topic employed immunohistochemical staining in an attempt to identify endothelial cells lacking staining in the lining of HHT-related arteriovenous malformations. ^33^ However, the *absence* of staining as a proxy for the *gain* of a mutation could be difficult to discover, especially if only a fraction of a cells would exhibit this lack of signal.

Using deep next-generation sequencing with unique molecular identifiers we successfully identified somatic mutations in multiple telangiectasia from different HHT patients. The somatic mutations we identified were present at frequencies ranging between 0.46% and 8.0% in the tissue, with an average of 2.3%. The low allele frequency is likely a result of two main contributing factors; the presence of normal tissue in the skin biopsy, and somatic mosaicism within the telangiectasia. The telangiectasias in this study were sampled as skin punch biopsies and although some of the surrounding skin tissue was removed before DNA extraction, an undetermined amount of normal tissue invariably remained. However, after removing the surrounding tissue, the enrichment for the vascular component of the tissue was subjectively greater than the often low somatic mutation allele frequency might suggest. We posit that a second explanation for the low mutant allele frequency is that telangiectasia are mosaic for the somatic mutation. This agrees with existing data from mouse models of HHT showing that induced retinal AVMs are mosaic: consisting of both heterozygous and homozygous null cells. ^34^ It is also consistent with the heterogeneity seen in Cerebral Cavernous Malformations ^35^ and the low mutant allele frequency in somatic mutations of other vascular malformation disorders. ^8–18^ Vascular malformations, similar to tumors in cancer, appear to be seeded by somatic mutations; however unlike tumors, vascular malformations do not appear to consist of pure populations of clonally expanded mutant cells, but contain a substantial percentage of unmutated cells.

In addition to the data presented here, the two-hit hypothesis of HHT-related vascular malformations is supported by evidence from mouse models of the disease. Whereas constitutional loss of both copies of *Eng* or *Acvrl1* in mice is embryonic lethal, mice heterozygous for constitutional deletion of either gene show extremely mild phenotypes with relatively few, if any, detectable vascular malformations. ^36,37^ Robust mouse models of HHT that recapitulate vascular malformation phenotypes require the use of Cre-Lox technology to delete both copies of *Eng* in a temporally controlled (postnatal), cell-type specific (endothelial cells) manner.

In these mouse models of HHT it is also required that biallelic KO of the HHT gene occur in endothelial cells. Several groups have experimented with expressing Cre recombinase in different vascular-related cell types including pericytes (NG2Cre), vascular smooth muscle cells (Myh11-Cre), and endothelial cells (Scl-Cre & Pdgfb-Cre); however, mice only develop vascular malformations when Cre is expressed in endothelial cells. ^38–41^ In addition, human *ENG* and *ACVRL1* are expressed primarily in endothelial cells, supporting that the somatic mutation must occur in an endothelial cell to seed a malformation. We have attempted to confirm that the somatic mutations we identified in human lesions occur in the endothelium by using laser capture microdissection, however these efforts have been hampered by the small quantity of tissue in telangiectasia biopsies and the difficulty of isolating a single layer of cells by microdissection. This question may be more easily addressed in larger arteriovenous malformations; however, these samples have been thus far inaccessible due to the rarity of their removal in HHT patients.

Interestingly, in addition to the local requirement for loss of both alleles, vascular malformations in this model only develop after injury such as an ear punch, or by VEGF injection. ^42^ This requirement of an angiogenic stimulus is consistent in mouse models for all HHT genotypes: *Eng*, *Acvrl1* and *Smad4*. ^43^ The requirement for knockout of both copies of the gene supports the genetic two-hit mechanism we describe here. In addition, the necessity for an angiogenic stimulus suggests that loss of both copies of the relevant HHT gene is necessary, but not sufficient, for the development of the vascular malformation.

We might have expected to find somatic mutations in every telangiectasia, however we only found somatic mutations in 9 of the 19 we sequenced. The next-generation sequencing strategy we employ for discovering somatic mutations is extremely sensitive for the detection of point mutations and small indels. However, there are several other types of genetic alterations that would result in biallelic loss of function due to loss of heterozygosity (LOH). LOH is a common occurrence in many tumors in cancer, and is a predominant mechanism of somatic loss/mutation. LOH can occur due to a variety of genetic mechanisms: large deletions, chromosome loss, and mitotic recombination. Given the apparent capacity for even a low fraction of somatically mutant cells to initiate the vascular malformation, it follows that the same would be true for LOH-associated mutational events; therefore, these mutations would appear instead as allelic imbalance rather than outright LOH. But if the level of allelic imbalance is as low as the frequency of somatic mutations we have observed in this study, we might expect linked marker haplotype ratios in the range of 48% to 52% at nearby markers; this slight and even trivial imbalance would be difficult if not impossible to detect and validate. It is also possible that non-genetic mechanisms such as loss of expression due to epigenetic silencing account for biallelic loss of function. This process, like the LOH associated events, would not be detected by our sequencing strategy; a problem that is only exacerbated by low allele frequency. Thus, it may not be surprising that we identified a somatic mutation in approximately only 19 of the VMs that were sequenced. We postulate it is highly likely that all of the 10 telangiectasia with no identified somatic mutation have biallelic loss by one of these other mechanisms.

One consequence of the genetic two-hit mechanism that might appear to be improbable is that, if true, a new somatic mutation must occur in every one of the numerous vascular malformations in HHT patients. For example, some HHT patients have dozens or more visible telangiectasia on the skin and mucocutaneous surfaces alone. ^44–46^ An attractive hypothesis to reconcile this conundrum would be if a somatic mutation first occurs in a circulating progenitor cell which then proliferates and seeds the formation of multiple telangiectasia. There is precedence for this mechanism in another vascular malformation syndrome, Blue Rubber Bleb Nevus syndrome. These patients display multiple small vascular lesions which harbor an identical somatic double-mutation in the *TIE2* gene. These vascular lesions appear to be anatomically-dispersed clones arising from an original, dominant, large lesion.9 In HHT-related vascular malformations, we report evidence that contradicts this hypothesis: different telangiectasia collected from the same individual harbor different, unique somatic mutations. This observation does not exclude the possibility that circulating cells may in some cases spread telangiectasia, however the data thus far suggests that the primary mechanism is independent somatic mutation events.

The dilemma of the requirement for numerous independent somatic mutation events in a single gene can be resolved with a probabilistic argument. Considering the size of the human genome (3.23• 10^9^ bp), the probability that a random somatic mutation occurs in the coding sequence of *ENG* (3201bp) in trans (50%with a pathogenic *ENG* germline mutation is 0.00005%. Compounding this value with empirical evidence that single cells have anywhere from 100-1500 somatic mutations per cell ^47–49^, and an estimate that 5.66% of exonic somatic mutations result in LOF ^47^; we calculate a conservative estimate that 0.00028% of cell have biallelic LOF *ENG* mutations. An adult human has at least 6• 10^11^ endothelial cells ^50^, therefore we estimate that an individual with HHT and a germline mutation in *ENG* has biallelic LOF *ENG* mutations in 1.5 million endothelial cells. It is clear that each cell with *ENG* biallelic LOF does not result in vascular malformation as individuals with HHT have at most hundreds, not millions, of telangiectasia. This disparity is consistent with the idea that telangiectasia only form under very specific conditions: likely that the somatic mutation must occur in a specific type of vascular bed, in an endothelial cell, and followed by local angiogenic stimulus.

Our samples consisted of telangiectasia from HHT patients from an HHT Centre of Excellence in Toronto, Canada. This group of patients had germline mutations in either *ENG* or *ACVRL1*, and somatic mutations were identified in telangiectasia from both genotypes. Mutations in *SMAD4* cause the combined syndrome HHT and Juvenile Polyposis (JP-HHT), however individuals with *SMAD4* mutations only account for 2% of HHT cases. ^3^ Unfortunately, no individuals with JP-HHT were present in our cohort, but we believe it is likely that JP-HHT-related telangiectasia follow an identical genetic two-hit mechanism, resulting from somatic mutations in *SMAD4*.

As HHT is caused by germline mutations in any one of 3 genes, an interesting question is whether the compound effect of a LOF mutation in two different causal genes could drive pathogenesis (e.g. a germline mutation in *ENG* and a somatic mutation in *ACVRL1*). However, thus far all of the somatic mutations we identified in HHT-related telangiectasia occur in the same gene as that harboring the germline mutation. This human genetic data is consistent with the observation that mice with combined deficiency for one allele each of *Acvrl1* and *Eng* are fertile, viable and are not teeming with vascular malformations ^51^ (Srinivasan and Marchuk, unpublished), as might be expected if trans-heterozygosity of mutations in these two HHT-genes could initiate vascular malformation development.

A genetic two-hit mechanism for HHT pathogenesis has therapeutic implications. Certain efforts to develop therapies for HHT assume a model of haploinsufficiency of the relevant HHT gene. These strategies attempt to increase the amount of the affected gene product by increasing the level of transcript/protein arising from the wild-type allele. ^52^ We show here that some fraction of cells within the malformation do not possess a wild-type allele. Even if expression in surrounding heterozygous cells could be increased, the null cells would remain devoid of protein, suggesting that this avenue of therapy may be ineffective. By contrast, a more effective strategy may be gene replacement, reintroducing a fully wild-type allele into the mutated cells ^53^, as this would simultaneously provide an extra copy of the gene to both the heterozygous and null cells. This strategy is particularly attractive as it might inhibit new VM formation by adding back a second wild-type copy of the mutated gene.

## Acknowledgments

We thank all patients and their families who participated in this study. We thank Dr. Nicolas Devos and the Duke Sequencing and Genomic Technologies Core for assistance designing experiments and sequencing. This study was supported by U.S. Department of Defense grant (W81XWH-17-1-0429) and a Fondation Leducq Transatlantic Network of Excellence Grant in Neurovascular Disease (17 CVD 03). MEF also received financial support from the Nelson Arthur Hyland Foundation.

